# A CHIMERIC RECEPTOR ENABLING ANTIBODY-GUIDED RETARGETING OF CAR T- CELLS

**DOI:** 10.64898/2026.04.27.720976

**Authors:** Natasha Vinanica, Jieying Yeo, Nurul Ain Nisrina Hamran, Dario Campana

## Abstract

Chimeric antigen receptor (CAR)-T cell therapies achieve remarkable responses in hematologic malignancies but are limited by target heterogeneity and antigen escape; therapeutic antibodies offer flexible targeting yet cannot leverage T-cell effector function. Integrating CAR-T cell potency with antibody targeting adaptability could expand the clinical reach of cellular immunotherapy. We engineered a CAR platform (termed CAR^FcR^) incorporating an Fc receptor (CD16^V158^) into the CAR extracellular domain, and enabling both CAR-mediated targeting and antibody-dependent recognition. CAR^FcR^-T cells were activated through either pathway, exhibiting robust cytokine secretion, degranulation, and proliferation. Anti-CD19 CAR^FcR^-T cells mediated cytotoxicity comparable to conventional CAR-T cells and eradicated CD19^+^ leukemia in xenograft models. Importantly, they also exerted antibody-dependent killing of CD19-negative tumor cells when guided by clinically approved antibodies, such as anti-HER2 trastuzumab and anti-CD20 rituximab. Dual engagement further enhanced cytotoxicity and enabled elimination of CD19-negative escape variants. This modular CAR^FcR^ architecture was successfully applied to additional CAR targets, including BCMA, CD123, and CD33, maintaining CAR function while allowing redirection with antibodies or CD16-binding bispecific engagers. Overall, CAR^FcR^ represents a versatile and clinically adaptable platform that integrates the strengths of CAR-T cells and antibody therapeutics to expand tumor targeting and overcome antigen escape.

## INTRODUCTION

Chimeric antigen receptor (CAR)-T cell therapy has demonstrated that genetically engineered lymphocytes can mediate profound and durable tumor regression.(1, 2) CAR engagement drives potent cytotoxic activation and sustained expansion, enabling high effector-to-target ratios in vivo. These properties have translated into remarkable clinical responses, particularly in hematologic malignancies.(3-11) However, the therapeutic success of CAR-T cells remains constrained by fixed antigen specificity, and relapse through antigen loss or modulation is a recurrent mechanism of resistance.(2, 12-15)

Monoclonal antibodies represent a cornerstone of cancer therapy. Although their tumor-targeting specificity parallels that of CARs, their anti-tumor activity is mediated through distinct effector mechanisms. A major component of their clinical efficacy derives from antibody-dependent cellular cytotoxicity (ADCC), which is initiated by engagement of Fcγ receptors on innate immune cells by the antibody Fc domain.(16, 17) Notably, therapeutic efficacy correlates with high-affinity Fcγ receptor polymorphisms, underscoring the importance of Fc-mediated effector engagement. (18-20) Yet T lymphocyte, the most potent and expandable cytotoxic effectors in adoptive cell therapy, lack activating Fcγ receptors and are therefore excluded from antibody-directed cytotoxic responses.(16, 17)

We hypothesized that integrating antibody-dependent recognition with CAR signaling in a single synthetic receptor would unify the proliferative and cytotoxic capacity of T cells with the targeting flexibility of monoclonal antibodies. To test this, we engineered chimeric receptors designed to enable post-infusion retargeting of adoptively transferred CAR-T cells. These receptors integrate two complementary modes of tumor recognition within a single molecular platform: direct antigen binding through an antibody-derived single-chain variable fragment (scFv), and indirect antigen recognition via high-affinity engagement of the Fc domain of tumor-bound antibodies.

## RESULTS

### A CD16-containing CAR confers antibody-binding capacity to T cells

To enable antibody-dependent targeting by CAR-T cells, we engineered a CAR incorporating the Fc-binding receptor CD16. The extracellular domain of CD16 containing the high-affinity V158 polymorphism was fused to the N-terminus of an anti-CD19-41BB-CD3ζ CAR via a flexible G4S1 linker (Fig. 1A). Peripheral blood T cells transduced with a retroviral vector encoding this construct (CAR^FcR^) showed robust surface expression of both the anti-CD19 single-chain variable fragment (scFv) and CD16 (Fig. 1 B-D). The proportion of scFv^+^ T cells generated with the CAR^FcR^ construct was comparable to that observed with a conventional anti-CD19 CAR (Fig. 1C and Fig. S1A). Because transduction efficiencies varied between donors, GFP encoded by the vector backbone was used as an internal reference. Correlation analyses between GFP and scFv expression demonstrated similar CAR expression levels in CAR^FcR^- and CAR-transduced T cells (Fig. S1B).

**Figure 1.**
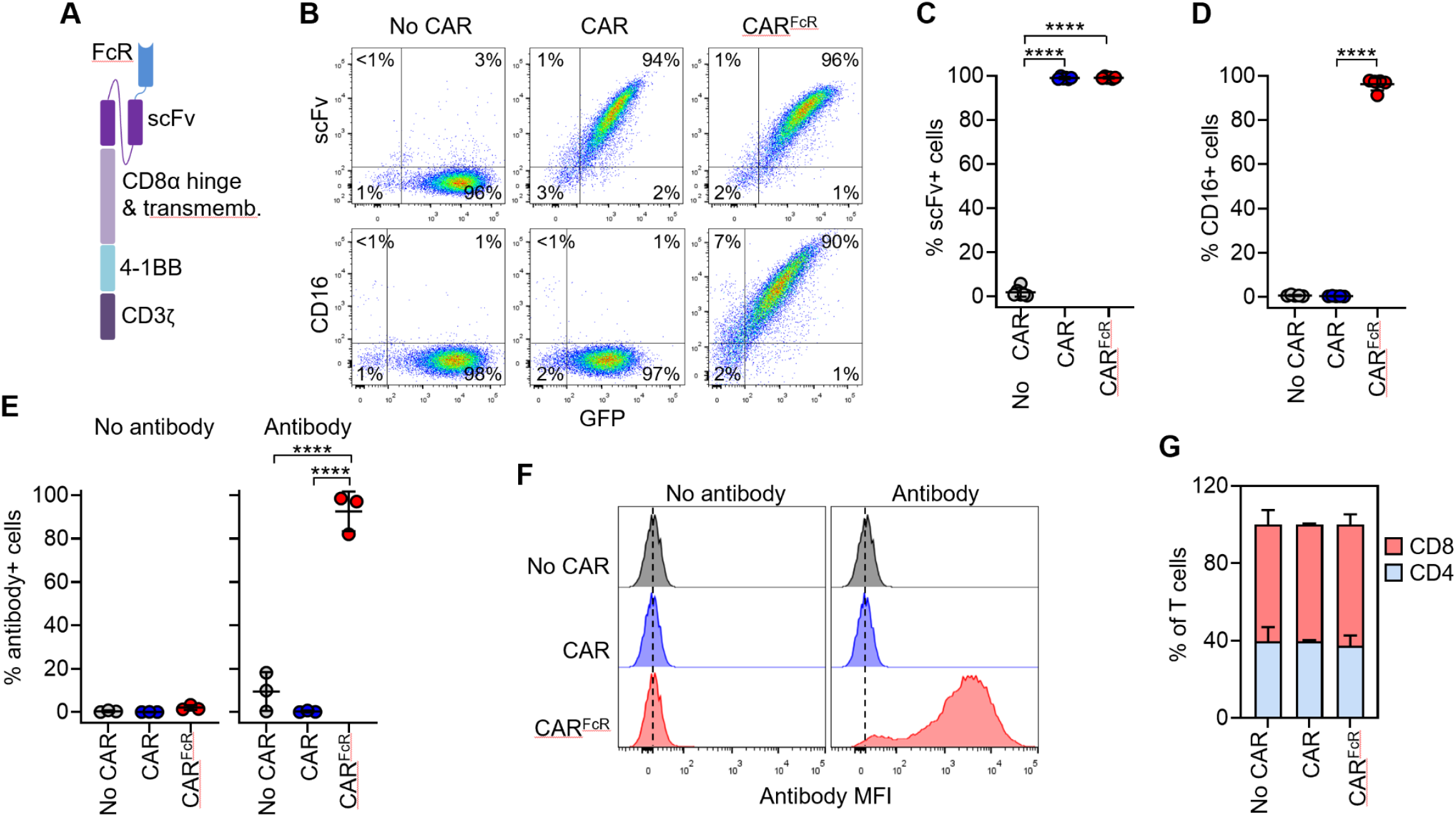
Design and expression of CAR^FcR^. (**A**) Schematic representation of CAR^FcR^, which includes an Fc receptor (FcR) and a single chain variable fragment (scFv) joined by a G4S1 linker (**B**) Flow cytometry plots show cell surface expression of anti-CD19 scFv (upper row; representative of six biological replicates) and of CD16 (lower row; representative of five biological replicates) on the Y-axes, compared to green fluorescence protein (GFP) levels on the X-axes. The percentage of cells in each quadrant is shown. (**C**) Percentage of GFP^+^ cells expressing anti-CD19 scFv shown in (B). Data are mean ± SD, with P values determined by unpaired t tests; n = 6 biological replicates. (**D**) Percentage of GFP^+^ cells expressing CD16 shown in (B). Data are mean ± SD, with P values determined by unpaired t tests; n = 5 biological replicates. (**E**) Percentage of antibody-bound cells after exposure to rituximab. Data are mean ± SD, with P values determined by unpaired t tests; n = 3 biological replicates. (**F**) Representative histograms of antibody-bound cells shown in (E). **(G)** Proportion of CD4+ and CD8+ T cells expressing anti-CD19 CAR^FcR^ or CAR. Data are mean ± SD from three biological replicates. ****P < 0.001.

We next tested whether CAR^FcR^ endowed T cells with the capacity to bind therapeutic antibodies. Following incubation with the anti-CD20 monoclonal antibody rituximab, antibody binding was readily detected on CAR^FcR^-T cells but not on conventional CAR-T cells (Fig. 1 E-F), confirming that the incorporated CD16 domain remained functionally accessible.

Introduction of the CD16 domain did not affect T-cell composition. CAR^FcR^- and conventional CAR-transduced products contained similar proportions of CD4^+^ and CD8^+^ T cells (Fig. 1G).

### CAR^FcR^ signaling is triggered by either antigen recognition or antibody engagement

Incorporation of CD16 into the CAR architecture did not impair signaling through the CD19-targeting scFv. To test this, anti-CD19 CAR^FcR^-T cells were co-cultured with the CD19^+^ acute lymphoblastic leukemia (ALL) cell line RS4;11. CAR^FcR^- and CAR-T cells showed comparable distributions of naive, central memory, effector memory, and effector subsets (Fig. 2A). CAR^FcR^-T cells upregulated the activation markers CD25 and CD69, produced interferon (IFN)-γ and tumor necrosis factor (TNF)-α, and underwent lytic granule exocytosis as measured by CD107a surface expression (Fig. 2 B-D). The magnitude of these responses was similar to that observed with conventional anti-CD19 CAR-T cells, indicating that addition of CD16 did not compromise CAR signaling.

**Figure 2.**
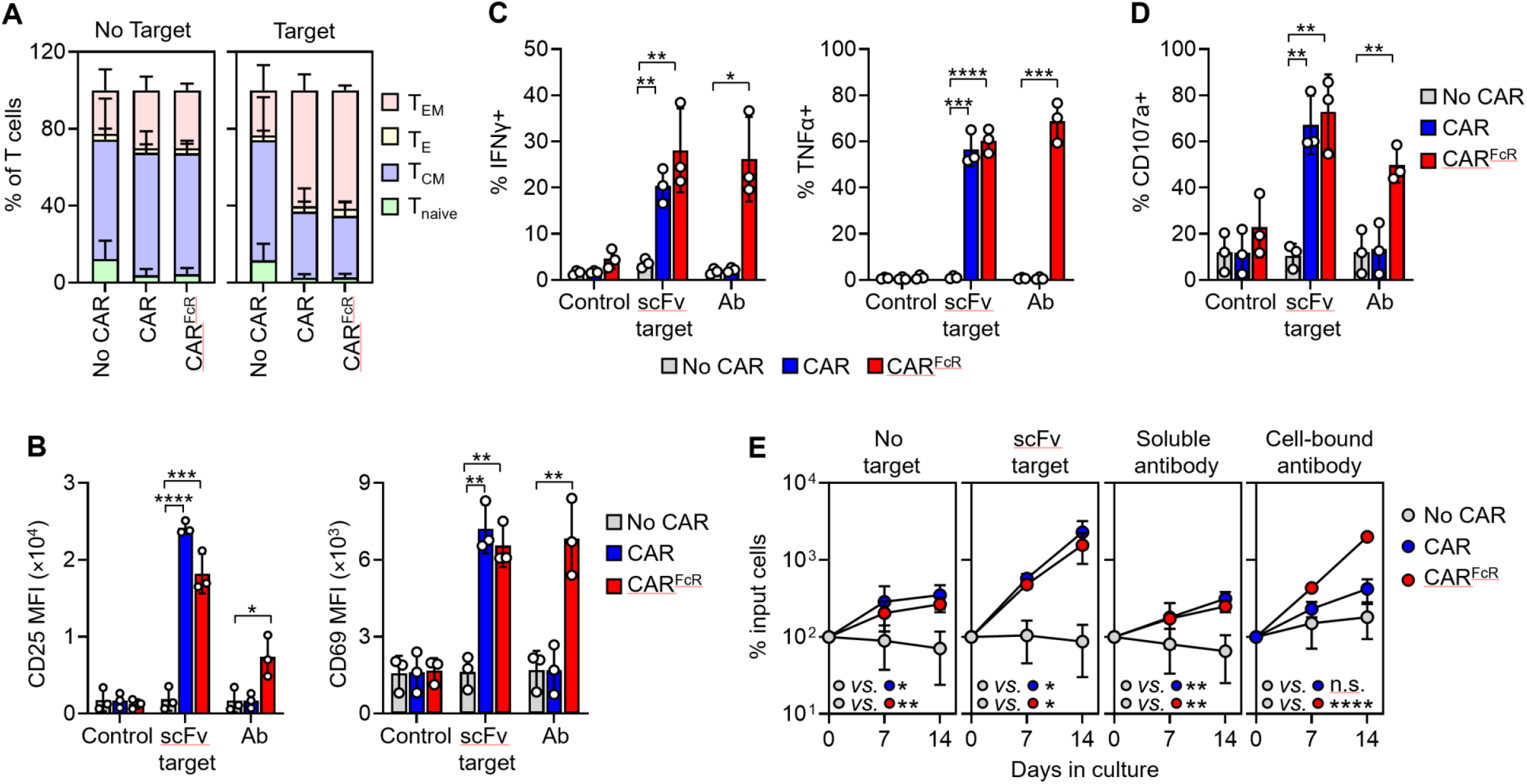
Signaling properties of CAR^FcR^. (**A**) Proportion of naive (T_naive;_ CD45RO-CD62L^+^), effector (T_E_; CD45RO-CD62L-), effector memory (T_EM_; CD45RO^+^ CD62L-), and central memory (T_CM_; CD45RO^+^ CD62L^+^) T cells expressing anti-CD19 CAR^FcR^ or CAR after 24-hour culture with or without target cells (RS4;11). (**B**) Expression of CD25 and CD69 in anti-CD19 CAR^FcR^ or CAR T cells after 24-hour culture with RS4;11 cells (scFv target) or surface-bound rituximab (Ab). (**C**) IFNγ and TNFα secretion after 6-hour stimulation of T cells as in (B). **(D)** Expression of CD107a in anti-CD19 CAR^FcR^- or CAR-T cells after 4-hour stimulation as in (B). (**E**) Proliferation of anti-CD19 CAR^FcR^- or CAR-T cells cultured alone, with irradiated RS4;11 cells, soluble antibody, or cell-bound antibody (rituximab-coated irradiated K562-CD20 cells) for 2 weeks. Each symbol indicates the percentage of cell recovery compared with the number of input cells. Data shown in (A) to (E) are mean ± SD from three biological replicates, with P values determined by unpaired t tests. *P < 0.05; **P < 0.01; ***P < 0.001; ****P < 0.0001.

We next asked whether engagement of the CD16 domain could independently trigger CAR^FcR^ activation. Cross-linking of CD16 using immobilized anti-CD20 antibody rituximab induced robust activation of CAR^FcR^-T cells, whereas conventional CAR-T cells remained inactive under these conditions (Fig. 2 B-D). These results indicate that CAR^FcR^ signaling can be initiated either by scFv-mediated recognition of CD19 or by antibody-dependent engagement of CD16.

CAR^FcR^ also retained the capacity to support antigen-driven T-cell expansion. When co-cultured with irradiated RS4;11 cells, anti-CD19 CAR^FcR^-T cells proliferated at rates comparable to those of conventional anti-CD19 CAR-T cells (Fig. 2E). In contrast, only CAR^FcR^-T cells expanded when cultured with irradiated K562-CD20 cells coated with rituximab. Soluble rituximab alone did not significantly affect T-cell proliferation (Fig. 2E).

### CAR^FcR^-T cells retain potent CD19-directed cytotoxic activity

We next evaluated whether incorporation of CD16 affected CAR-mediated cytotoxic activity. Short-term (4-hour) cytotoxicity assays were performed using a panel of CD19^+^ ALL cell lines (RS4;11, Nalm-6, OP-1, and REH) and CD19^+^ lymphoma cell lines (Ramos and Daudi). In head-to-head comparisons, anti-CD19 CAR^FcR^-T cells displayed cytotoxic activity comparable to that of conventional anti-CD19 CAR-T cells across all target cell lines tested (Fig. 3A and Fig. S2). Longer-term cytotoxicity assays using Nalm-6 cells as targets yielded similar results, confirming that incorporation of CD16 did not impair the capacity of the CAR to mediate target cell killing (Fig. 3B).

**Figure 3.**
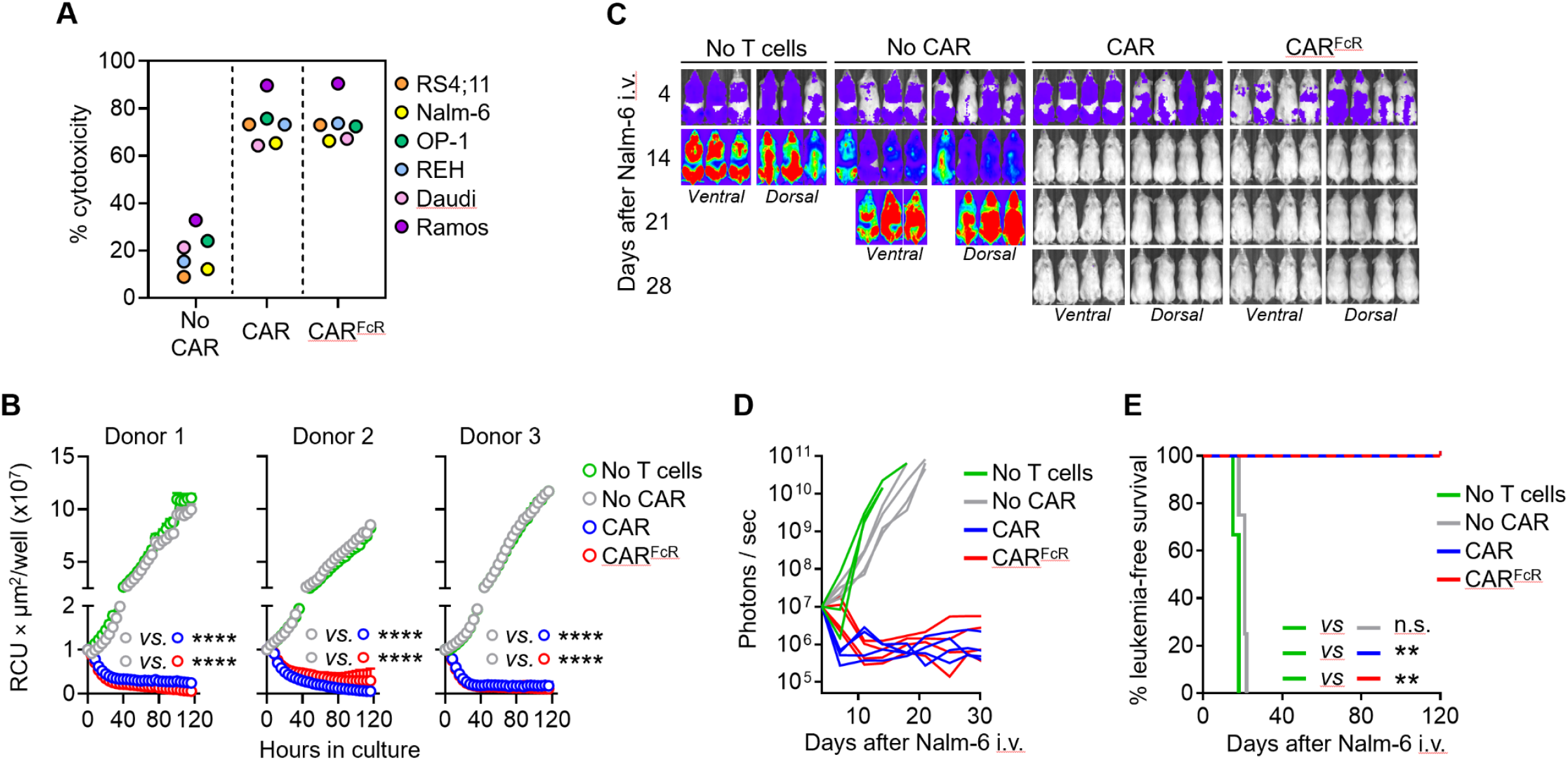
Anti-CD19 CAR^FcR^-T cells exert CAR-directed cytotoxicity. (**A**) Cytotoxicity of anti-CD19 CAR^FcR^- and CAR-T cells against CD19^+^ ALL and lymphoma cell lines at a 1:1 E:T ratio in 4-hour assays. Each symbol indicates mean of three biological replicates with each cell line. The full set of data for each cell line is shown in Fig. S2. (**B**) Cytotoxicity of anti-CD19 CAR^FcR^- and CAR-T cells against mCherry-expressing Nalm-6 cells at a 1:2 E:T ratio monitored by real-time Incucyte imaging for 5 days. Each symbol represents mean ± SD of technical triplicates at the indicated time points, with P values determined by unpaired t tests. Data were normalized to the first time point. (**C**) NSG mice were injected with luciferase-expressing Nalm-6 cells (0.5 × 10^6^ IV) and treated with T cells expressing GFP only (no CAR), anti-CD19 CAR, or CAR^FcR^ (20 × 10^6^ IV) four days later; a control group was left untreated left (no T cells). Images on day 4 were taken with enhanced sensitivity to better detect tumor engraftment. The full set of images is shown in Fig. S3. (**D** and **E**) Luminescence measurements (D) and Kaplan-Meier survival curves (E) of mice shown in (C). The full data set for (C) and (D) is shown in Fig. S3. **P < 0.01; ****P < 0.0001; n.s., not significant.

To determine whether these findings translated in vivo, we used a xenograft model in which NOD-scid-IL2RG^null^ (NSG) mice were engrafted with luciferase-expressing Nalm-6 leukemia cells and subsequently treated with either anti-CD19 CAR^FcR^-T cells or conventional anti-CD19 CAR-T cells. Treatment with either CAR-T cell product eradicated leukemia and resulted in durable leukemia-free survival (Fig. 3 C-E and Fig. S3 A-B). In contrast, Nalm-6 leukemia cells rapidly expanded in untreated mice and in mice receiving control T cells lacking CAR expression. Analysis of peripheral blood parameters showed no significant differences in platelet counts, red blood cell counts, or hemoglobin levels between mice treated with CAR^FcR^- or conventional CAR-T cells (Fig. S3 C-E).

### Antibody-dependent targeting broadens CAR^FcR^ tumor recognition

Because CAR^FcR^ signaling can be triggered by engagement of antibody Fc domains, we reasoned that CAR^FcR^-T cells might kill tumor cells that lack the CAR cognate antigen if those cells are coated with a therapeutic antibody. To test this hypothesis, we co-cultured anti-CD19 CAR^FcR^-T cells with the HER2^+^ breast cancer cell line SKBR3, which does not express CD19. Robust killing of SKBR3 cells was observed when tumor cells were coated with the anti-HER2 monoclonal antibody trastuzumab (Fig. 4A and Fig. S4). In contrast, SKBR3 cells expanded in the absence of trastuzumab, or when CAR^FcR^-T cells were replaced with conventional anti-CD19 CAR-T cells, regardless of whether trastuzumab was present (Fig. 4A and Fig. S4).

**Figure 4.**
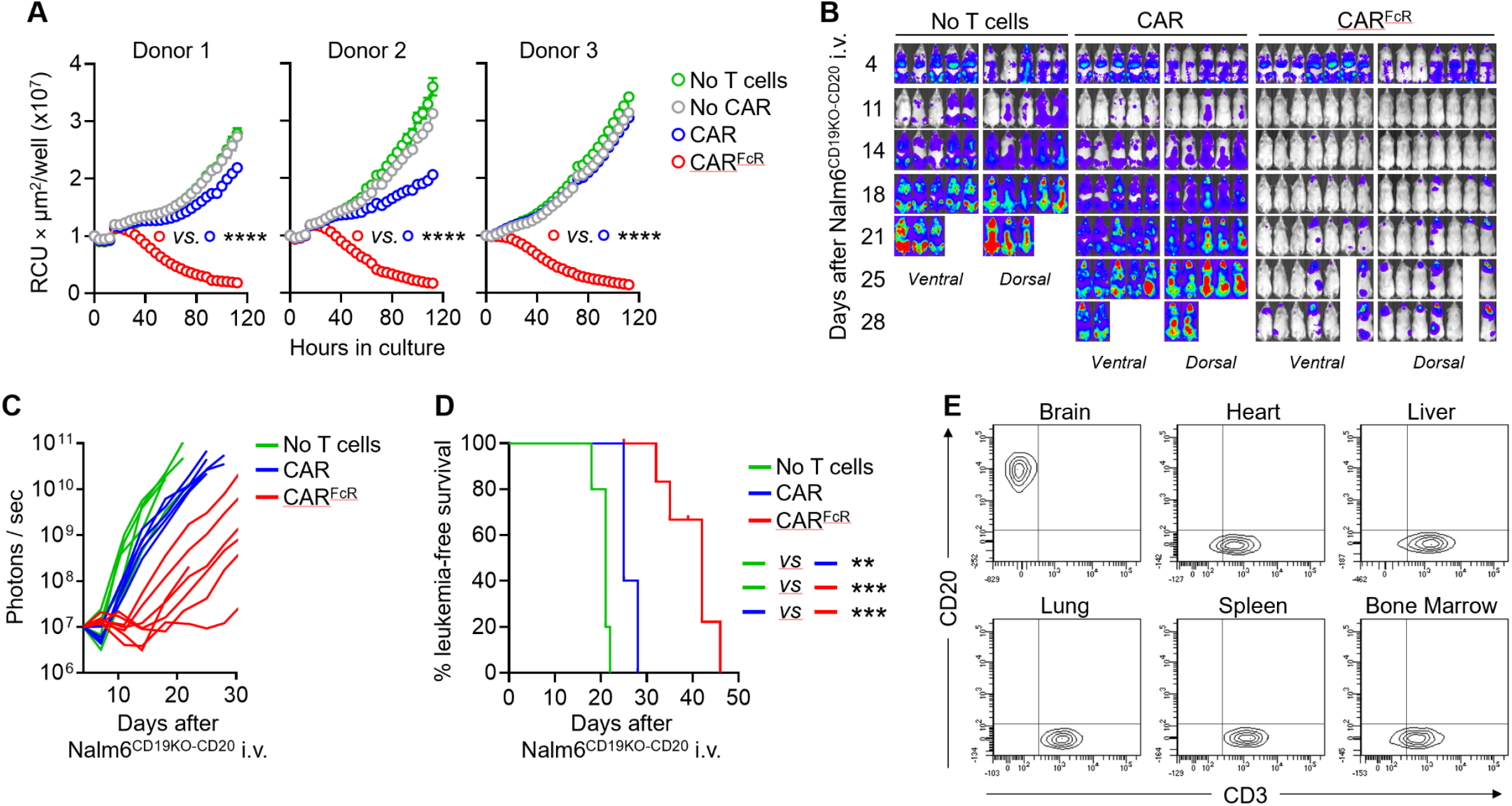
Anti-CD19 CAR^FcR^-T cells can kill target cells by antibody-dependent cell cytotoxicity (ADCC). (**A**) ADCC of anti-CD19 CAR^FcR^-T cells against mCherry-expressing HER2^+^ SKBR3 cells in the presence of trastuzumab at a 1:2 E:T ratio, monitored by real-time Incucyte imaging for 5 days. Each symbol represents mean ± SD of technical triplicate at the indicated time points, with P values determined by unpaired t tests. Data were normalized to the first time point after addition of T cells to the cultures. The results of parallel cultures without trastuzumab are shown in Fig. S4. (**B**) NSG mice were injected with luciferase-expressing Nalm6^CD19KO-CD20^ cells (0.5 × 10^6^ IV) followed by rituximab alone (10 mg/kg IP) or in combination with CAR^FcR^- or anti-CD19 CAR-T cells (20 × 10^6^ IV) four days later. Mice images on day 4 were taken with enhanced sensitivity to detect tumor engraftment. The full set of images is shown in Fig. S5. (**C**) Luminescence measurements of mice shown in (B). **(D)** Kaplan-Meier survival curves of mice shown in (B). (**E**) Flow cytometry plots show Nalm6^CD19KO-CD20^ cells (CD20^+^CD3^−^) and T cells (CD3^+^) in the organs harvested from a mouse with recurrent disease. The full data set for (C) to (E) is shown in Fig. S5. Data for two additional mice are shown in Fig. S6. **P < 0.01; ***P < 0.001 ****P < 0.0001.

We next evaluated ADCC mediated by CAR^FcR^-T cells in vivo. NSG mice were engrafted with luciferase-expressing Nalm-6 leukemia cells in which CD19 expression had been ablated by CRISPR-Cas9 and CD20 was introduced by retroviral transduction (Nalm6^CD19KO-CD20^). Mice were then treated with the anti-CD20 antibody rituximab together with either anti-CD19 CAR^FcR^-T cells or conventional anti-CD19 CAR-T cells. Treatment with CAR^FcR^-T cells plus rituximab produced a marked reduction in tumor burden and sustained suppression of tumor growth (Fig. 4B and Fig. S5A). In contrast, minimal antitumor activity was observed in mice treated with rituximab alone or with rituximab plus conventional anti-CD19 CAR-T cells. Consistent with these findings, mice receiving the combination of CAR^FcR^-T cells and rituximab showed significantly lower bioluminescence signals (Fig. 4C and Fig. S5 B-C) and prolonged survival compared with the other treatment groups (Fig. 4D and Fig. S5D).

Despite the initial therapeutic response, signs of relapse emerged approximately three weeks after infusion of Nalm6^CD19KO-CD20^ cells, particularly in the cranial region. To investigate the cause of relapse, three mice with recurrent disease were analyzed after euthanasia. High frequencies of CD20^+^CD3− cells, consistent with Nalm6^CD19KO-CD20^ leukemia cells, were detected in the brain but not in other organs (Fig. 4E and Fig. S6). In contrast, human T cells (CD3^+^) were present in most peripheral tissues but were detected in the brain of only one of the three mice. This distribution suggests that limited penetration of monoclonal antibodies across the blood-brain barrier may have restricted effective tumor targeting in the central nervous system, allowing leukemia relapse at this site.

### CAR^FcR^ enables T-cell cytotoxicity against multiple targets simultaneously or sequentially

Because CAR^FcR^ supports dual modes of antigen recognition, through the CAR scFv or through CD16 engagement by antibody-coated targets, we examined how simultaneous engagement of both pathways affects T-cell function. To this end, we assessed cytotoxicity against the CD19^+^CD20^+^ Daudi lymphoma cell line. Anti-CD19 CAR^FcR^-T cells and conventional anti-CD19 CAR-T cells showed comparable cytotoxicity against Daudi cells under baseline conditions (Fig. 5A). However, when Daudi cells were coated with the anti-CD20 antibody rituximab, cytotoxicity mediated by CAR^FcR^-T cells was significantly increased, whereas cytotoxicity mediated by conventional CAR-T cells remained unchanged (Fig. 5A).

**Figure 5.**
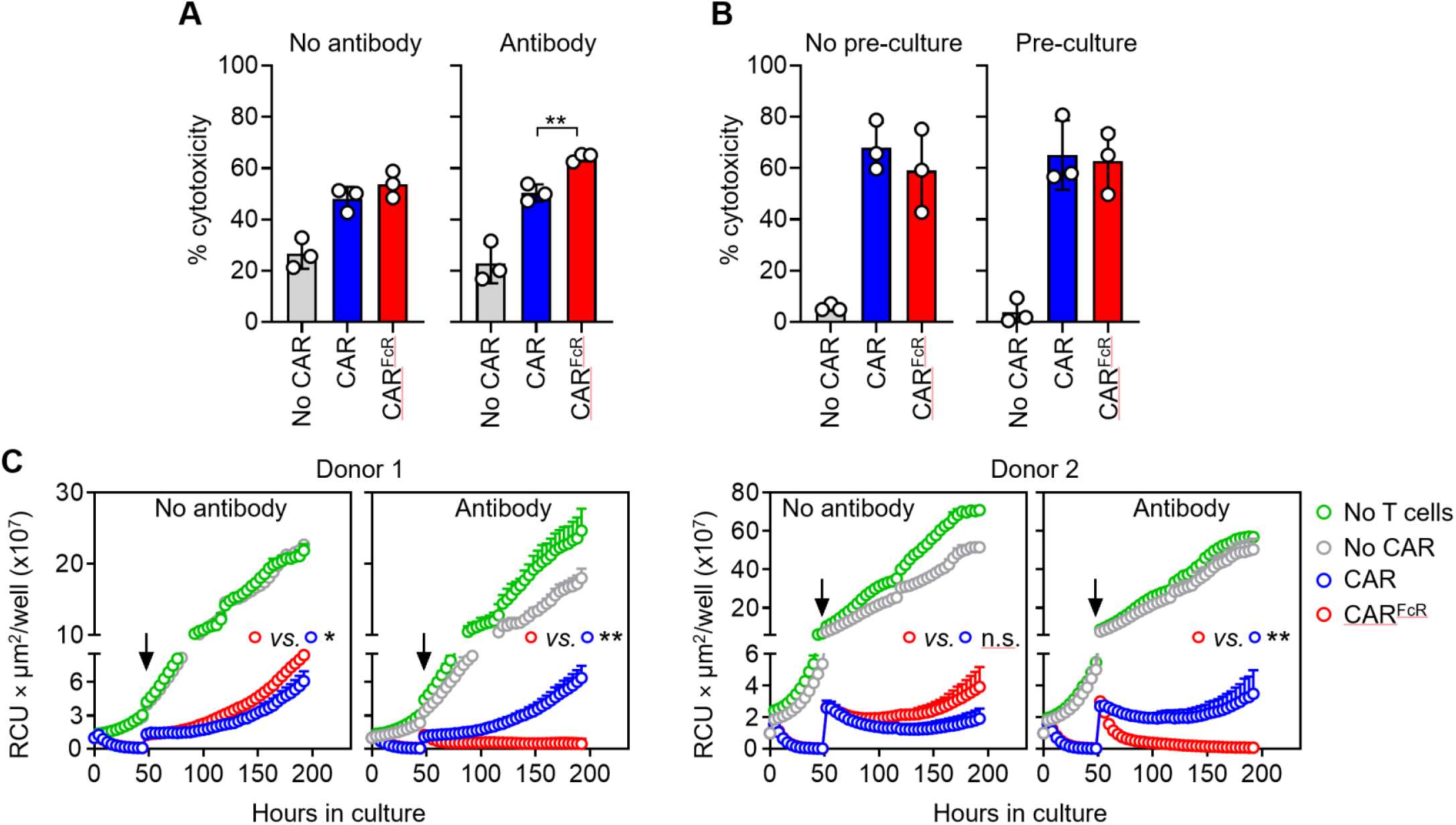
Anti-CD19 CAR^FcR^-T cells can target multiple antigens simultaneously or sequentially. (**A**) Cytotoxicity of anti-CD19 CAR^FcR^- and CAR-T cells against CD19^+^CD20^+^ Daudi cells at a 1:2 E:T ratio in 4-hour assays, with or without rituximab. Data shown are mean ± SD, with P values determined by unpaired t tests; n = 3 biological replicates. (**B**) Cytotoxicity of anti-CD19 CAR^FcR^- and CAR-T cells against RS4;11 cells in 4-hour assays, after pre-culture with rituximab-coated CD20-expressing K562 cells for 24 hours. Data shown are mean ± SD, with P values analyzed by unpaired t tests; n = 3 biological replicates. (**C**) Cytotoxicity of anti-CD19 CAR^FcR^- and CAR-T cells against Nalm6^CD19KO-CD20^ cells after co-culture with wild type Nalm-6 at a 1:1 E:T ratio for 2 days. Both target cells express mCherry and their growth was monitored by real-time Incucyte imaging for 8 days. Arrow indicates the time of Nalm6^CD19KO-CD20^ cells addition to the cultures. Data were normalized to the first time point. Each symbol represents mean ± SD of technical triplicates at the indicated time points, with P values determined by unpaired t tests. *P < 0.05; **P < 0.01; n.s., not significant.

We also tested whether antibody engagement of CD16 might interfere with CAR-mediated function. Anti-CD19 CAR^FcR^- and conventional CAR-T cells were first co-cultured with rituximab-coated K562 cells genetically modified to express CD20 (K562-CD20) and subsequently rechallenged with CD19^+^ RS4;11 leukemia cells. Cytotoxic activity against RS4;11 cells was comparable regardless of prior exposure to rituximab-coated targets (Fig. 5B), indicating that CD16 engagement does not impair CAR signaling and may instead augment cytotoxic responses.

Loss of target antigen expression is a well-recognized mechanism of tumor escape following CAR-T cell therapy.(2, 12-15) To determine whether CAR^FcR^-T cells could overcome antigen loss through antibody-mediated redirection, we modeled CD19-negative relapse in vitro. Anti-CD19 CAR^FcR^- or conventional CAR-T cells were first co-cultured with wild-type Nalm-6 cells until the CD19^+^ population was eliminated, after which the T cells were rechallenged with Nalm6^CD19KO-CD20^ cells. Conventional anti-CD19 CAR-T cells were unable to control the outgrowth of Nalm6^CD19KO-CD20^ cells (Fig. 5C). In contrast, CAR^FcR^-T cells completely suppressed their expansion in the presence of rituximab (Fig. 5C), demonstrating that antibody-mediated targeting can redirect CAR^FcR^ cytotoxicity toward antigen-negative tumor variants.

### The CAR^FcR^ platform can be extended to multiple targets and antibody-based engagers

We evaluated whether the CAR^FcR^ architecture could be further optimized by modifying the position of the CD16 domain or the linker length (Fig. S7A). Four alternative CAR^FcR^ designs were generated and expressed in peripheral blood T cells (Fig. S7 B-C).

Comparison of their capacity to induce T-cell activation (Fig. S7 D-E), ADCC (Fig. S7F), and CAR-mediated cytotoxicity (Fig. S7G and S8) revealed that none of these alternative configurations presented clear functional advantages over the original construct used in the experiments described above, indicating that the baseline CAR^FcR^ design provides robust activity.

We next assessed whether the functional properties of CAR^FcR^ were restricted to the anti-CD19 CAR or could be generalized to other tumor targets. To this end, CAR^FcR^ constructs incorporating scFvs directed against B-cell maturation antigen (BCMA), CD123 or CD33 were generated. Retroviral transduction of T cells resulted in high surface expression of both the scFv and CD16 for all three constructs (Fig. 6A).

**Figure 6.**
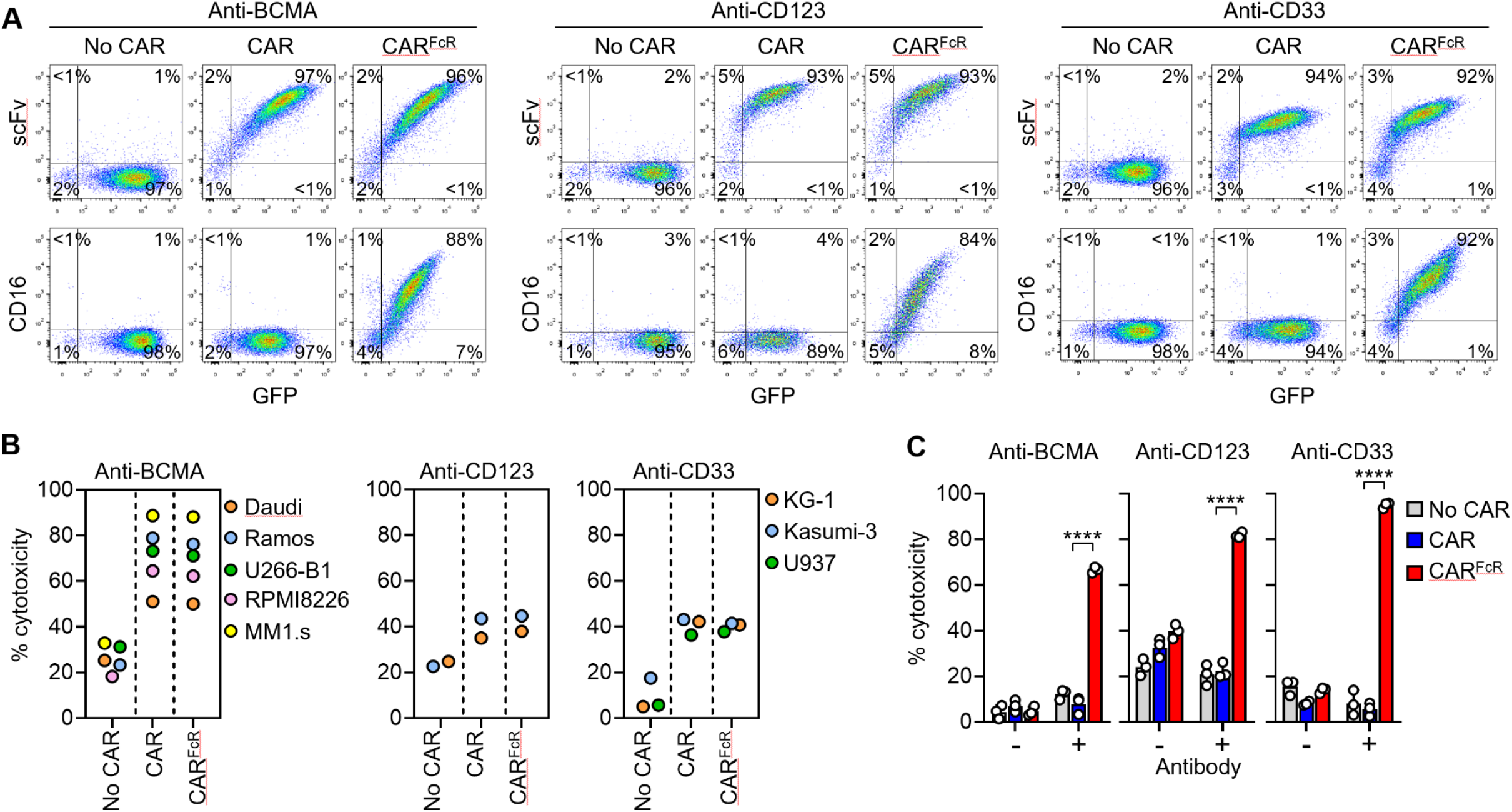
The CAR^FcR^ architecture can be adapted to CAR antigens beyond CD19. (**A**) Flow cytometry plots show surface expression of anti-BCMA, anti-CD123 and anti-CD33 CAR scFvs (upper row), and of CD16 (lower row) on T cells transduced with the corresponding CAR^FcR^ and parental CAR constructs. The percentage of cells in each quadrant is shown. (**B**) Cytotoxicity of anti-BCMA CAR^FcR^, anti-CD123 CAR^FcR^, and anti-CD33 CAR^FcR^- and CAR-T cells against their respective target cells at a 1:1 E:T ratio in 4-hour assays. Each symbol indicates mean of technical triplicate with each cell line; the full set of data is shown in Fig. S9. (**C**) ADCC of anti-BCMA CAR^FcR^, anti-CD123 CAR^FcR^, and anti-CD33 CAR^FcR^- and CAR-T cells against HER2^+^ SKBR3 with or without trastuzumab at a 4:1 E:T ratio. Data are shown as mean ± SD, with P values analyzed by unpaired t tests; n = 3 technical replicates. ****P < 0.0001.

In 4-hour cytotoxicity assays against BCMA^+^ lymphoma cell lines Daudi and Ramos, and multiple myeloma cell lines RPMI8226, MM1.s, and U266-B1, anti-BCMA CAR^FcR^-T cells displayed potent cytotoxic activity comparable to that mediated by the parental anti-BCMA CAR lacking CD16 (Fig. 6B and Fig. S9). Similarly, anti-CD123 CAR^FcR^-T cells were cytotoxic against CD123^+^ acute myeloid leukemia (AML) cell lines KG-1 and Kasumi-3, and cytotoxicity was comparable to that of T cells expressing anti-CD123 CAR without CD16 (Fig. 6B and Fig. S9). Finally, anti-CD33 CAR^FcR^- and conventional anti-CD33 CAR-T cells exhibited comparable cytotoxicity against CD33^+^ AML cell lines KG-1, Kasumi-3, and U937 (Fig. 6B and Fig. S9).

These findings indicate that incorporation of CD16 into the CAR architecture does not impair CAR-mediated cytotoxicity across a range of target antigens. Conversely, the capacity of CAR^FcR^-T cells to mediate ADCC was independent of scFv specificity. T cells expressing anti-BCMA, anti-CD123, or anti-CD33 CAR^FcR^ constructs efficiently killed HER2^+^ SKBR3 cells in the presence of the anti-HER2 antibody trastuzumab, whereas no cytotoxicity was observed with the corresponding CAR-T cells lacking CD16 (Fig. 6C).

In addition to clinically approved antibodies, CD16 expression also enables CAR^FcR^-T cells to be redirected using engineered bispecific engagers containing CD16-binding domains. To test this approach, we generated a bispecific engager composed of an anti-CD16 scFv fused to an anti-CD33 scFv (21) and added it to co-cultures of anti-CD19 CAR^FcR^-T cells with CD33^+^ KG-1 and Kasumi-3 AML cells. The engager induced robust cytotoxic activity by anti-CD19 CAR^FcR^-T cells against CD33^+^ targets, whereas no effect was observed when CAR^FcR^-T cells were replaced with conventional anti-CD19 CAR-T cells (Fig. 7A).

**Figure. 7.**
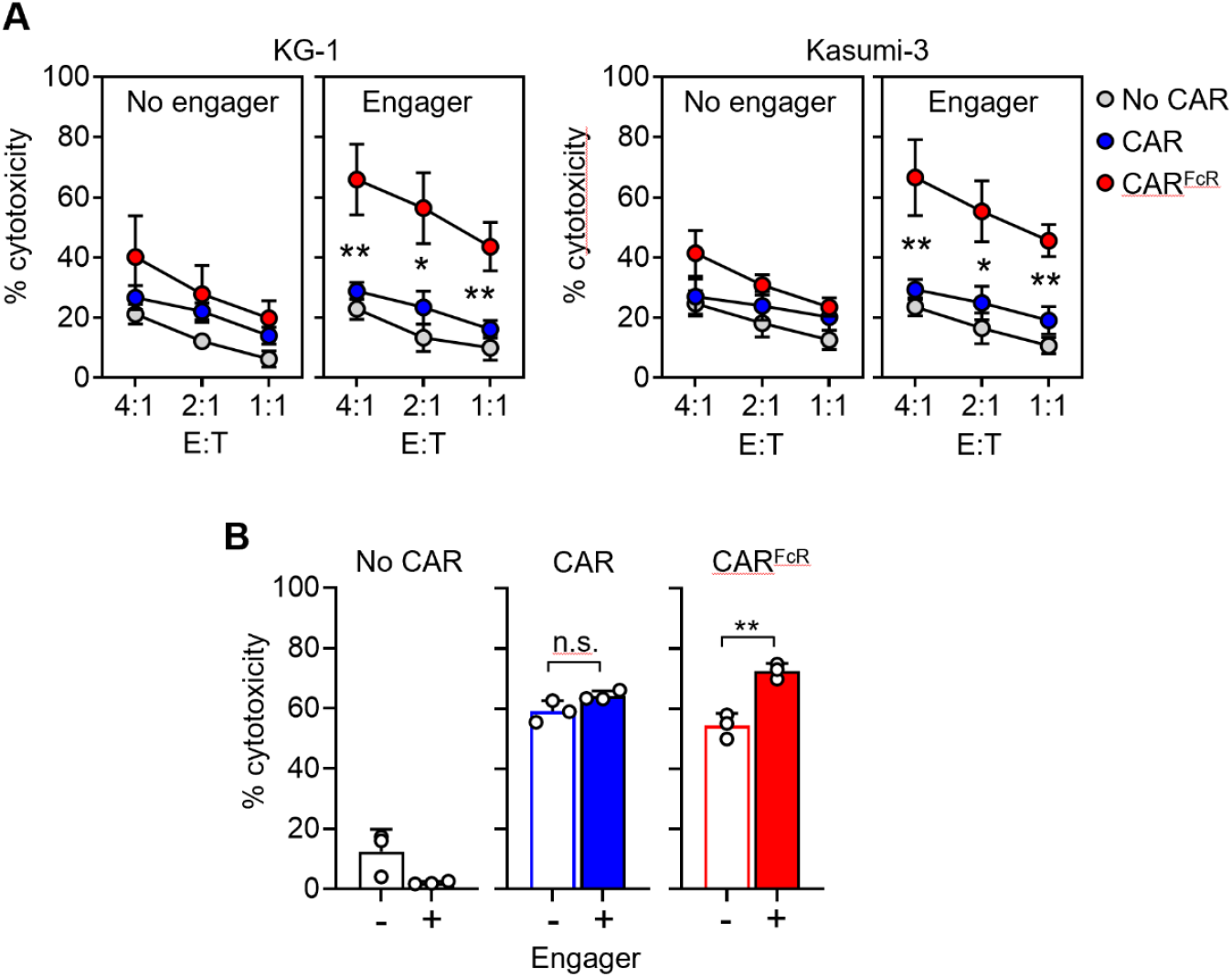
CAR^FcR^-T cells can be stimulated with an anti-CD16 bispecific engager. (**A**) Cytotoxicity of anti-CD19 CAR^FcR^- and CAR-T cells against CD33^+^ KG-1 and Kasumi-3 cells with or without a bispecific engager containing scFvs against CD16 and CD33 in 4-hour assays at the indicated E:T ratios. Data are shown as mean ± SD, with P values determined by unpaired t tests; n = 3 biological replicates. (**B**) Cytotoxicity of anti-CD123 CAR^FcR^-T cells against CD123^+^CD33^+^ KG-1 cells with or without the bispecific engager as in (A) in 4-hour assays at 2:1 E:T ratio. Data shown are mean ± SD, with P values determined by unpaired t tests; n = 3 technical replicates. n.s, not significant; *P < 0.05; **P < 0.01.

Moreover, enhanced target cell killing was observed when anti-CD123 CAR^FcR^-T cells were co-cultured with CD123^+^CD33^+^ KG-1 cells in the presence of the engager (Fig. 7B).

Together, these findings demonstrate that the CAR^FcR^ platform can be broadly applied across different CAR specificities and can be redirected toward additional tumor antigens using either therapeutic antibodies or engineered bispecific engagers.

## DISCUSSION

This study describes a CAR platform that integrates conventional CAR-mediated cytotoxicity with ADCC. By incorporating the high-affinity Fc receptor CD16^V158^ into an anti-CD19-4-1BB-CD3ζ CAR backbone, we generated chimeric receptors (CAR^FcR^) capable of recognizing tumor antigens through the CAR scFv while simultaneously engaging therapeutic antibodies through Fc binding. CAR^FcR^ constructs were robustly expressed in human T cells and retained dual functionality, enabling both direct CAR-mediated killing and antibody-guided cytotoxicity. Importantly, this architecture was not restricted to CD19 targeting: CAR^FcR^ constructs incorporating scFvs directed against BCMA, CD123, or CD33, were efficiently expressed and supported both CAR-driven cytotoxicity and antibody-mediated redirection. Together, these findings establish CAR^FcR^ as a modular platform that expands the functional scope of CAR-T cells by enabling antibody-guided engagement of additional tumor antigens.

The CAR^FcR^ design preserves conventional CAR activity while enabling CAR-T cells to be redirected by antibodies against additional antigens. This creates post-infusion flexibility without new cell manufacturing or repeat lymphodepletion. This feature could be particularly valuable for addressing antigen escape, a major driver of relapse after CD19-directed CAR-T therapy in B-cell malignancies.(2, 12-15) Persistent CAR^FcR^-T cells could be redirected with clinically available antibodies against alternative B-cell antigens such as CD20, CD22, or CD38. Multi-antigen targeting could also be implemented from the outset to counter antigen expression heterogeneity, or insufficient CAR reactivity against cells with low antigen density.(22) Consistent with this concept, CAR^FcR^-T cells mediated CAR-dependent cytotoxicity and ADCC both concurrently and sequentially. Extending this approach to targets such as CD33 or CD123(23-26) could similarly help address antigen heterogeneity in AML.(27, 28)

ADCC is a key mechanism of action for many therapeutic antibodies and is typically initiated through engagement of CD16A (FCGR3A) on natural killer (NK) cells by the Fc domain of antibodies bound to tumor cell surfaces.(16, 17) Patients with the FCGR3A V158 polymorphism have improved responses to antibody-based therapies, consistent with the higher affinity of this variant for IgG immune complexes.(18-20) Importantly, CD16 functions as a low-affinity Fc receptor that preferentially recognizes immune complexes or antibodies immobilized on cell surfaces, while interacting minimally with monomeric circulating IgG.(17, 29) This property favors selective activation in the presence of antibody-opsonized tumor cells. In line with this notion, CAR^FcR^-T cells remained inert in the presence of soluble antibodies but were potently activated by antibodies immobilized on target cells. Adoptive transfer of T cells engineered to express Fc-guided chimeric receptors alone has previously been tested clinically with rituximab in refractory B-cell NHL, demonstrating clinical responses, long-term persistence, and limited toxicity.(30-32) Our approach extends this concept by integrating Fc receptor function directly into a conventional CAR framework. Most approved CAR-T cells products incorporate second-generation 4-1BB (CD137) costimulatory domains as in our CAR^FcR^ construct;(33-35) whether other costimulatory molecules could be advantageous in the CAR^FcR^ context remains to be determined.(36) Antibody opsonization of tumor cells should increase effective target density for CAR^FcR^, broadening tumor cell recognition and/or enhancing signaling. It remains unclear whether these receptors behave as Boolean “OR” receptors,(37) where engagement of a single binding domain elicits a maximal signal equivalent to dual engagement, or whether simultaneous engagement of both the scFv and Fc receptor domains produces additive or cooperative increases in signaling strength.

Current approaches to multi-antigen targeting include infusing multiple CAR-T products or co-expressing several CARs within a single T cell, strategies that increase manufacturing complexity and can impair receptor expression or function.(38) A central objective of this study was therefore to develop a receptor that would confer ADCC gain-of-function without compromising the expression or activity of the original CAR. In the selected constructs, CAR-mediated cytotoxicity and proliferation were largely indistinguishable from those of the parental CAR. Compared with alternative designs such as dual-scFv CARs or other multi-target receptors,(38-40) CAR^FcR^ offers distinct advantages: antigen specificity can be determined through antibody selection, new targets can be introduced after CAR-T infusion, and activity can be dynamically regulated through antibody dosing. Other groups have developed CAR systems activated by antibody-linked adaptors or bispecific engagers,(41, 42) but these strategies rely on engineered reagents that are not routinely used clinically and may not fully recapitulate the effector functions of conventional therapeutic antibodies.

CAR-T therapy has transformed the treatment of B-cell malignancies but remains limited by antigen heterogeneity, clonal escape and the difficulty of reproducing optimal signaling across different CAR targets.(1, 2) The CAR^FcR^ platform addresses these challenges by enabling flexible multi-antigen targeting while preserving the functional integrity of well-validated CAR designs. In addition, it creates opportunities to combine T-cell cytotoxicity with established mechanisms of antibody therapy, including complement activation and recruitment of innate immune effector cells. In sum, the CAR^FcR^ architecture enables dynamic, multi-antigen targeting using clinically approved antibodies and provides a practical strategy to mitigate antigen escape and tumor heterogeneity without re-engineering the cellular product.

## MATERIALS AND METHODS

The objective of this study was to develop a chimeric receptor that integrated antibody-dependent antigen recognition with CAR signaling. The receptor, termed CAR^FcR^, was generated by fusing the extracellular domain of the Fc receptor CD16 to a CAR. The activation, cytokine secretion, proliferation, and cytotoxicity of T cells upon CAR^FcR^ engagement were studied in vitro and compared to those elicited by CARs lacking CD16.

The antitumor capacity of CAR^FcR^-T cells was also studied in murine xenograft models, where mice in different treatment groups were randomized based on bioluminescence signals after tumor cell engraftment to ensure balanced tumor burden; investigators were not blinded to the mouse groupings. Sample sizes for animal experiments were determined based on prior experience with these animal models; no animal was excluded due to illness from the study.

All animal experiments were approved by the Institutional Animal Care and Use Committee of the National University of Singapore. Immortalized human cell lines and primary human T cells were used in this study. Primary human T cells were isolated from discarded tubing from platelet donations obtained from the Health Sciences Authority Blood Bank (Singapore), with approval from the Institutional Review Board of the National University of Singapore. Details of the reagents and methods used are provided in Supplementary Materials.

Statistical analyses were performed using GraphPad Prism version 10.6.1. Statistical tests and the experimental replicates are indicated in figure legends. Data are presented as individual values or as mean ± SD from at least three independent experiments. Comparisons of two groups were performed using two-tailed unpaired t test unless otherwise indicated.

Survival was analyzed using Kaplan-Meier curves, and probability (P) values were calculated using the log-rank (Mantel-Cox) test. P-values are denoted with asterisks as follows: not significant (n.s.), P > 0.05; *P < 0.05; **P < 0.01; ***P < 0.001; and ****P < 0.0001.

## Authors’ contributions

N.V. designed constructs, performed experiments, analyzed data; J.Y and N.A.N.H. performed experiments; D.C. initiated the study and analyzed data. N.V. and D.C drafted the manuscript, which all authors have reviewed.

