## Supplementary Methods and Supplementary Figures S1 to S9 for "A CHIMERIC RECEPTOR ENABLING ANTIBODY-GUIDED RETARGETING OF CAR T- CELLS"

**Supplementary Material:**

Supplementary Methods

Supplementary Figures S1 – S9

### SUPPLEMENTARY METHODS

#### Cells and culture conditions

The cell lines RS4;11, REH and Nalm-6 from B-cell acute lymphoblastic leukemia (ALL); Daudi and Ramos from B-cell non-Hodgkin lymphoma (NHL); KG-1 and Kasumi-3 from acute myeloid leukemia (AML); K562 from chronic myelogenous leukemia (CML); U937 from histiocytic lymphoma; U266-B1, RPMI8226, and MM1.s from multiple myeloma; and SKBR3 from breast cancer were obtained from the American Type Culture Collection (ATCC; Rockville, MD). The B-ALL cell line OP-1 was previously developed in our laboratory.(33) The genes encoding mCherry, firefly luciferase, CD20 or anti-CD16 scFv-anti-CD33 scFv bispecific engager(21) were expressed using a murine stem cell virus (MSCV) retroviral vector (from St. Jude Children's Research Hospital, Memphis, TN). Cell lines were maintained in RPMI-1640 (HyClone, Marlborough, MA) supplemented with 10-20% fetal bovine serum (FBS) (HyClone) and 1% penicillin-streptomycin (Gibco, Waltham, MA).

CD19-negative Nalm-6 cells expressing CD20 (Nalm6$^{CD19KO-CD20}$) were generated by electroporating CRISPR-CD19 gRNA (Genscript, Piscataway, NJ) targeting exon 3 into wild type Nalm-6 using Amaxa Cell Line Nucleofector Kit V (Lonza, Basel, Switzerland) and the Nucleofector II program C-005 (Lonza). The CRISPR-CD19 gRNA-electroporated Nalm-6 was subsequently sorted using a MoFlo Astrios cell sorter (Beckman Coulter, Brea, CA) to obtain pure population of CD19-negative Nalm-6 cells, which were then transduced to express CD20.

Peripheral blood mononuclear cells were isolated by density gradient centrifugation from discarded anonymized byproducts of platelet donations from the Health Sciences Authority Blood Bank (Singapore) with the approval of the Institutional Review Board of the National University of Singapore. T cells were expanded from peripheral blood mononuclear cells using TransAct (Miltenyi Biotec, Bergisch Gladbach, Germany) and cultured in RPMI-1640 supplemented with 10% FBS and 1% penicillin-streptomycin. Interleukin-2 (IL-2; Proleukin, Novartis, Basel, Switzerland) was added at 120 IU/mL every 2 - 3 days.

**Gene cloning and retroviral transduction**

The anti-CD19-41BB-CD3ζ CAR was previously made in our laboratory.(33) The extracellular domain of CD16 with V158 polymorphism was fused to N-terminus of anti-CD19 single-chain variable fragment (scFv, clone FMC63) with a G4S1 linker (Fig. 1A). A similar approach was used to generate the variants with different position of CD16 domain and linker lengths (Fig. S7A). Receptors targeting BCMA, CD123, or CD33 were generated using scFvs derived from antibodies against BCMA (clone C11D5.3), CD123 (clone 26292), or CD33 (clone hP67.6). All constructs were subcloned into an MSCV-IRES-GFP vector between EcoRI and XhoI restriction sites.

For retroviral transduction, MSCV retroviral vector-conditioned medium was added to RetroNectin (Takara, Otsu, Japan)-coated polypropylene tubes; after centrifugation and removal of the supernatant, T cells were added to the tubes and left at 37°C for 24 hours; fresh viral supernatant was added again after 24 hours. After 48 hours, the transduced T cells were washed twice and maintained in RPMI-1640 with 10% FBS, 1% penicillin-streptomycin, and 200 IU/mL IL-2 until the time of the experiments, 7-21 days after transduction.

**Detection of surface receptor expression and target binding**

Surface expression of CAR scFv was detected using a biotin-conjugated goat anti-mouse $F(ab')_2$ antibody followed by streptavidin conjugated to APC (both from Jackson

ImmunoResearch, West Grove, PA). Surface expression of CD16 was detected using anti-CD16-APC antibody (clone B73.1, BD Biosciences, Franklin Lakes, NJ).

Antibody-binding capacity of CD16 was assessed by exposing T cells to 1 μg/mL of anti-CD20 antibody rituximab (Selleckchem, Houston, TX) followed by goat anti-human IgG conjugated to PE (Southern Biotech, Birmingham, AL).

Cell staining was analyzed using Fortessa flow cytometer (BD Bioscience), with BDFACS Diva (BD Biosciences) or FlowJo software (FlowJo, Ashland, OR).

**Immunophenotyping, cytotoxicity, and proliferation**

T-cell immunophenotyping was performed with anti-CD3 APC (clone SK7; BD Biosciences), anti-CD4 V450 (clone RPA-T4; BD Biosciences), and anti-CD8 PE (clone RPA-T8; BD Biosciences). In some experiments, T cells were cultured with RS4;11 cells at a 1:1 E:T ratio for 24 hours before staining with anti-CD45RO-PE/Cy7 (clone UCHL; BD Pharmingen) and anti-CD62L-APC (clone DREG-56; BD Pharmingen).

To assess CAR-mediated cytotoxicity, target cells labeled with calcein red-orange AM (Thermo Fisher Scientific, Waltham, MA) were co-cultured with T cells for 4 hours at the indicated E:T ratios. In some experiments, 1 μg/mL anti-CD20 antibody rituximab (Selleckchem) or 15 ng/mL anti-CD16-anti-CD33 bispecific engager were added to the cultures. Viable target cells were counted using the Accuri C6 Plus flow cytometer (BD Biosciences).

To measure antibody-dependent cellular cytotoxicity (ADCC), T cells were co-cultured with $HER2^+$ SKBR3 cells expressing luciferase at the indicated E:T ratios in the presence of 10 μg/mL anti-HER2 antibody trastuzumab (Selleckchem). Bright-Glo luciferase reagent (Promega, Madison, WI) was added 4 hours after the co-culture. Luminescence signal was measured 5 minutes later using BioTek FLx800 plate reader (BioTek, Winooski, VT) and analyzed with BioTek Gen5 2.0 data analysis software.

Long-term cytotoxicity was performed by co-culturing T cells with mCherry-expressing target cells at the indicated E:T ratios, with the addition of 100 IU/mL IL-2 every 2 -3 days. Viable mCherry$^+$ cells were monitored with real-time Incucyte Zoom System (Essen BioScience, Ann Arbor, MI), acquiring whole-well images every 4 hours, with cell number expressed as red calibrated units (RCU) x $\mu m^2$/well.

To assess sequential cytotoxicity, T cells were first co-cultured with CD20-expressing K562 (K562-CD20) cells at a 1:1 E:T ratio in the presence of 1 µg/mL rituximab for 24 hours to induce ADCC. The cells were then transferred to RS4;11 cells labelled with Far Red (Thermo Fisher Scientific) and incubated for an additional 4 hours to evaluate CAR-mediated cytotoxicity.

In other experiments, T cells were first co-cultured with wild type Nalm-6 expressing mCherry for 2 days at 1:1 E:T ratio before re-challenged with Nalm6$^{CD19KO-CD20}$ expressing mCherry at 1:1:1 ratio in the presence or absence of 1 µg/mL rituximab. Viable mCherry$^+$ cells were monitored with real-time Incucyte imaging, as described above.

To measure T-cell proliferation, T cells were cultured alone or in the presence of irradiated (100 Gy) target cells at a 1:1 ratio, with the addition of 100 IU/mL IL-2 every 2 -3 days. Irradiated target cells were added at the beginning of the culture and every 7 days thereafter. In parallel experiments, 1 µg/mL rituximab was added to the culture once a week. The number of viable GFP-positive T cells was counted with the Accuri C6 Plus flow cytometer.

**Cell activation, degranulation, and cytokine production**

To measure T-cell activation, T cells were cultured alone or with target cells for 24 hours. Expression levels of activation markers were measured using anti-CD25-PE (clone 2A3, BD Biosciences) and anti-CD69-APC (clone L78, BD Biosciences). In some experiments, activation marker expression was assessed after culturing T cells in rituximab-coated plates for 24 hours. To prepare the plate, 100 μL of 1 μg/mL rituximab was added to 96-well flat-bottom plates (Costar, Corning, NY) and incubated at 4°C overnight. The plates were washed once with 1 x DPBS (Gibco) before use.

Exocytosis of lytic granules was assessed by measuring CD107a expression after culturing T cells with target cells at a 1:1 ratio. In some experiments, T cells were cultured in rituximab-coated plates prepared as described above. Anti-human CD107a-PE (clone H4A3; BD Biosciences) was added at the beginning of the culture. After 1 hour, monensin (BD GolgiStop) was added, and the cells were cultured for an additional 3 hours before flow cytometric analysis.

To measure TNFα and IFNγ cytokine secretion, T cells were cultured with target cells at a 1:1 ratio or in rituximab-coated plates, as described above. After 1 hour, brefeldin A (BD GolgiPlug) was added and the cultures were continued for another 5 hours. The cells were then stained intracellularly with anti-TNFα-APC (clone 6401.1111; BD Biosciences) or anti-IFNγ-PE (clone 25723.11; BD Biosciences) and analyzed on a flow cytometer.

**Xenograft models**

To assess anti-leukemic activity in vivo, 0.5 x $10^6$ luciferase-expressing Nalm-6 cells were injected intravenously (IV) into NOD.Cg-Prkd$^{cscid}$ IL2rg$^{tm1Wjl}$/SzJ (NOD-scid-IL2RG$^{null}$, NSG; Jackson Laboratory, Bar Harbor, ME) mice. After tumor cells had engrafted four days later, mice were assigned to four groups with equivalent tumor loads and received IV injections of 20 x $10^6$ T cells expressing GFP, CAR, or CAR$^{FcR}$. One group received culture medium only as control.

In a second model, mice were injected IV with 0.5 x $10^6$ luciferase-expressing Nalm6$^{CD19KO-CD20}$ cells. After four days, the mice received IV injections of 20 x $10^6$ T cells expressing either anti-CD19 CAR or CAR$^{FcR}$, along with weekly intraperitoneal (IP) injections of rituximab (10 mg/kg; Teva Pharmaceutical, Tel Aviv, Israel). In both models, all mice received 20,000 IU of IL-2 IP every 2-3 days.

Tumor progression was monitored by measuring the luminescence signal with a Xenogen IVIS-200 System (PerkinElmer, Waltham, MA), following IP injection of aqueous D-luciferin potassium salt (150 µg/g body weight; PerkinElmer). The luminescence signal was analyzed with Living Image 3.0 software. Mice were euthanized when luminescence reached 1 x $10^{11}$ photons/second or when physical signs warranted euthanasia.

Blood was collected via cheek prick, and cell counts were determined with a Celltac α hematology analyzer (Nihon Kohden, Tokyo, Japan). After treatment with red blood cell lysis solution (MilliporeSigma, Burlington, MA), the cells were stained with anti-human CD45-APC (clone 2D1; BioLegend), anti-human CD3-PE/Cy7 (clone HI30; BD Pharmingen), and anti-mouse CD45-PE (clone 30-F11; BD Pharmingen).

Selected organs (brain, heart, liver, lung, and spleen) and tissue (bone marrow) were collected from mice in the second mouse model following euthanasia for flow cytometric analysis. Organs were minced and passed through a 40 µm cell strainer (Corning), then filtered through a 30 µm pre-separation filter (Miltenyi Biotec) before treatment with red blood cell lysis solution. Cells were subsequently stained with anti-human CD20-APC (clone L27; BD Biosciences), anti-human CD3-PE/Cy7, and anti-mouse CD45-PE.

All procedures were approved by the Institutional Animal Care and Use Committee of the National University of Singapore.

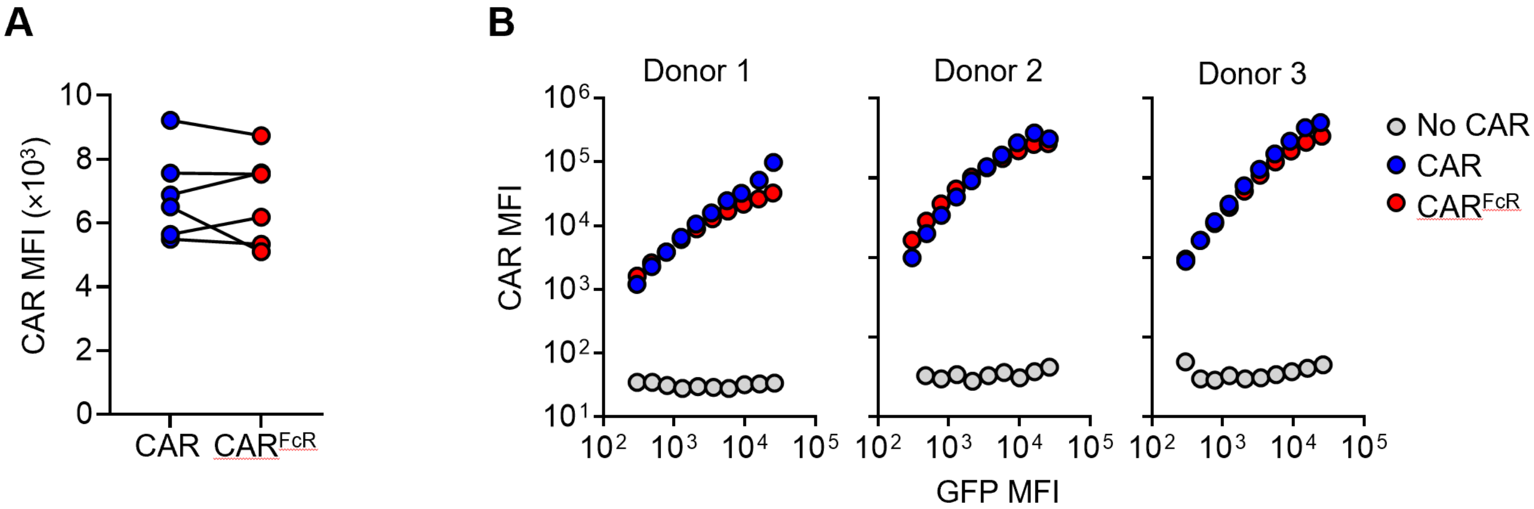


**Figure S1. Expression of anti-CD19 CAR$^{FcR}$ in primary T cells.** (**A**) Graph shows mean fluorescence intensity (MFI) of T cells transduced with anti-CD19 CAR$^{FcR}$ or anti-CD19 CAR. Data from six biological replicates are shown. (**B**) Relation between MFI of CAR scFv and MFI of GFP of T cells transduced with GFP only (no CAR), anti-CD19 CAR or anti-CD19 CAR$^{FcR}$. Each symbol corresponds to measurement at the CAR MFI at the indicated level of GFP expression.

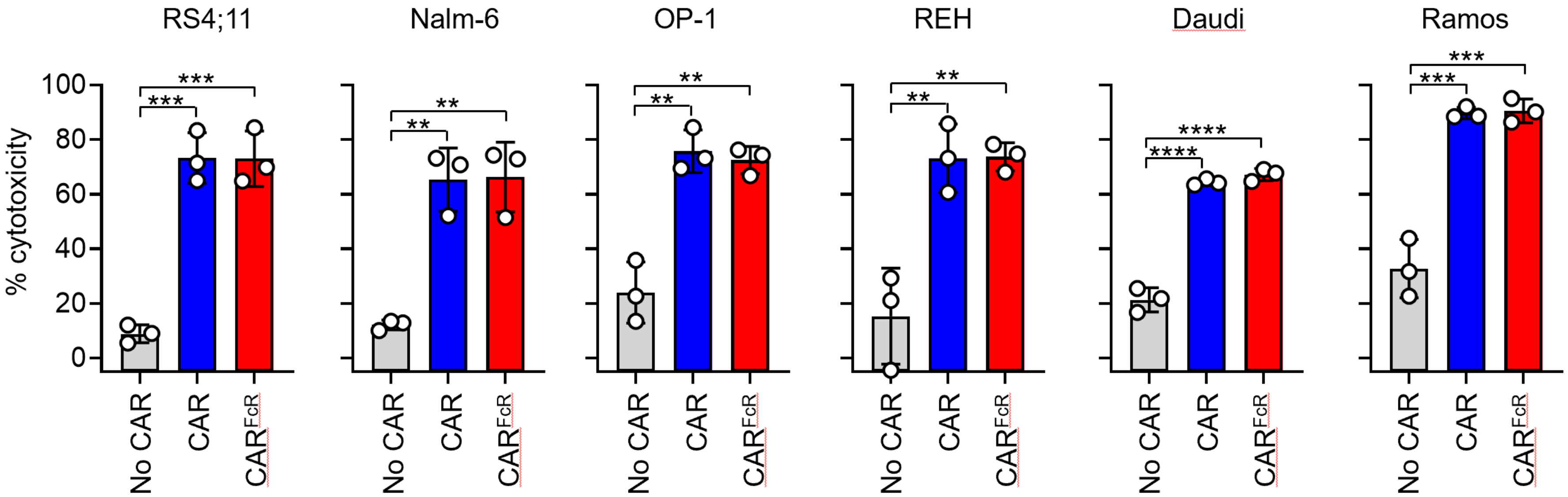


**Figure S2. CAR-mediated cytotoxicity of anti-CD19 CAR$^{FcR}$-T cells against CD19$^{+}$ cell lines.** Cytotoxicity of T cells transduced with GFP only (no CAR), anti-CD19 CAR or anti-CD19 CAR$^{FcR}$ against the CD19$^{+}$ ALL cell lines RS4;11, Nalm-6, OP-1, and REH, or the lymphoma cell lines Daudi and Ramos at a 1:1 E:T ratio in 4-hour assays. Data shown are mean ± SD, with P values determined by unpaired t tests; n = 3 biological replicates. **P < 0.01; ***P < 0.001; ****P < 0.0001.

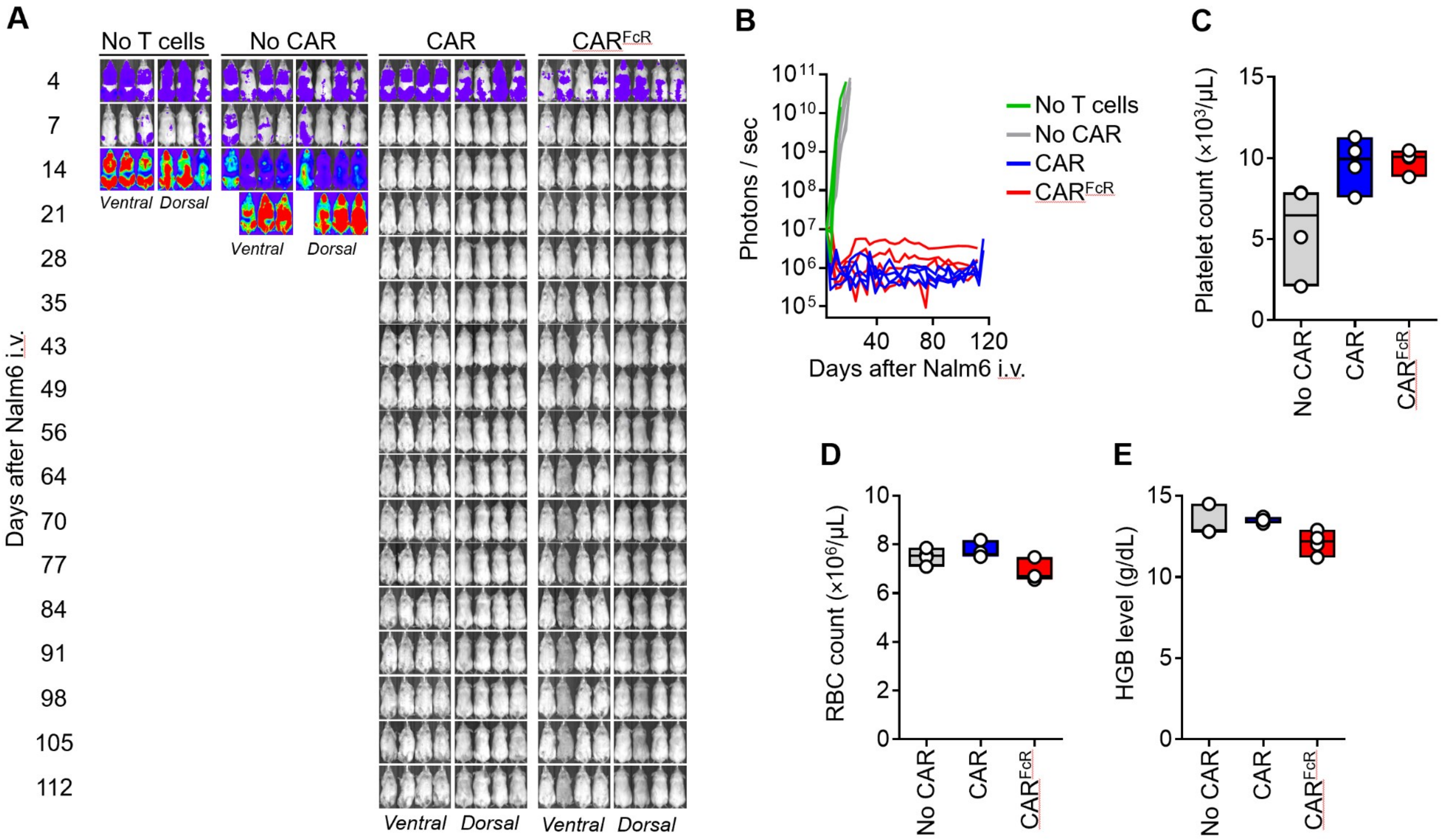


**Figure S3. Anti-tumor activity of anti-CD19 CAR$^{FcR}$-T cells in a xenograft model of ALL.** (**A**) NSG mice were injected IV with 0.5 x $10^6$ Nalm-6 cells that had been transduced with luciferase; 4 days later, 20 x $10^6$ T cells expressing GFP only (no CAR), anti-CD19 CAR or anti-

CD19 CAR$^{FcR}$ were administered by IV injection. Ventral and dorsal images show Nalm-6 cells engraftment as measured by luminescence after IP injection of aqueous d-luciferin potassium salt (150 µg/g body weight). The figure shows the complete set of images; images on day 4 were taken with enhanced sensitivity to better detect tumor engraftment. Images from days 4-28 are also shown in Fig 3C. (**B**) Luminescence measurements in the groups of mice shown in (A). (**C)** Platelet counts, (**D**) red blood cell (RBC) counts, and (**E**) hemoglobin (HGB) levels in the mouse blood 14 days after T-cell infusion. Data in (C) to (E) are presented as box plots, with dots indicating measurement from one mouse (n = 4 for all groups) and lines indicating the median.

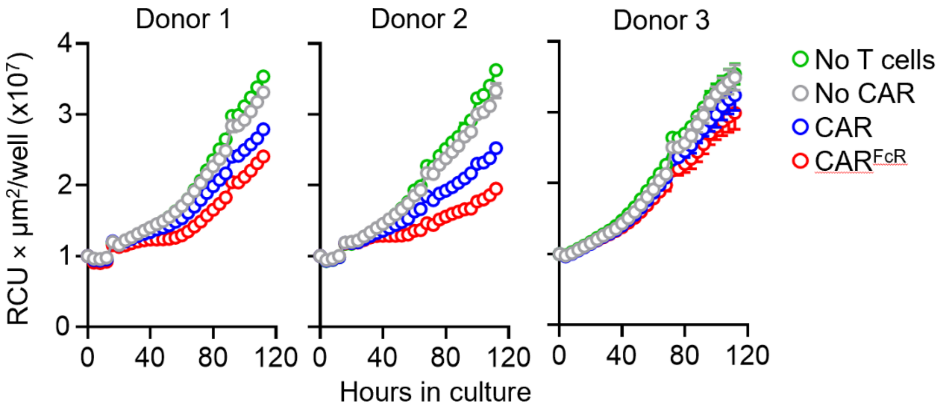


**Figure S4. Antibody is required to trigger antibody-dependent cell cytotoxicity (ADCC) by anti-CD19 CAR$^{FcR}$-T cells.** T cells expressing GFP only (no CAR), anti-CD19 CAR, or anti-CD19 CAR$^{FcR}$ were cultured with mCherry-expressing HER2$^{+}$ SKBR3 cells at a 1:2 E:T ratio in the absence of trastuzumab and monitored by real-time Incucyte imaging for 5 days. The results of parallel cultures with trastuzumab are shown in Fig. 4A. Each symbol represents the mean ± SD of technical triplicate at the indicated time points. Data were normalized to the first time point after the addition of T cells to the cultures.

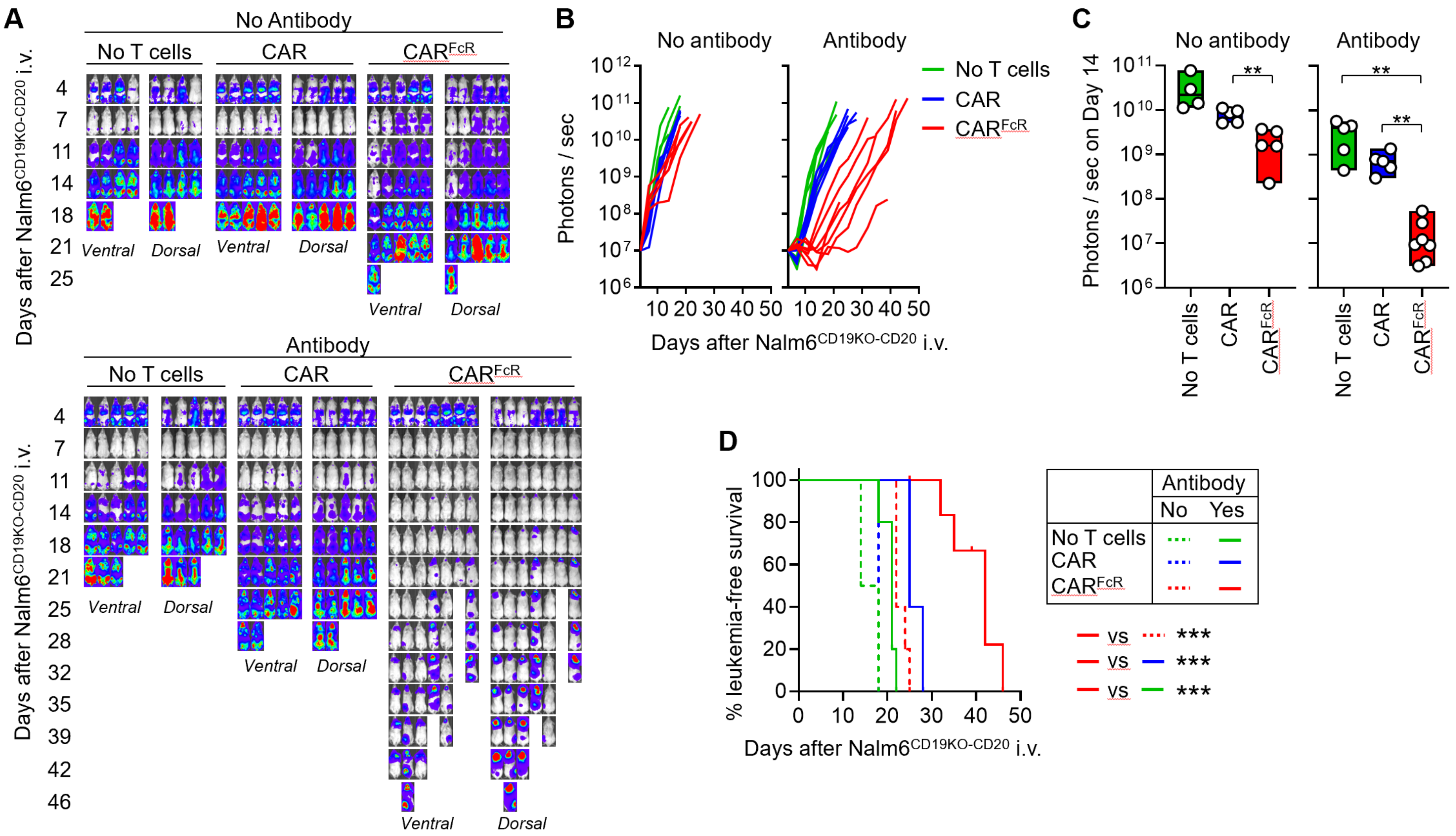


**Figure S5. Anti-CD19 CAR$^{FcR}$-T cells exert antibody-dependent cell cytotoxicity (ADCC) in vivo.** (**A**) Luciferase-expressing Nalm6$^{CD19KO-CD20}$ cells (0.5 x $10^6$) were injected IV in NSG mice, followed by IP injection of rituximab alone (10 mg/kg) or in combination with 20 x $10^6$ T cells expressing either anti-CD19 CAR or anti-CD19 CAR$^{FcR}$ four days later. Ventral and dorsal images show tumor cell engraftment as measured by luminescence after IP injection of aqueous d-luciferin potassium salt (150 µg/g body weight). Images on day 4 were taken with enhanced sensitivity to detect tumor engraftment. The figure shows the complete set of images; images of mice treated with antibody taken on days 4-28 are also shown in Fig. 4B. (**B**) Luminescence measurements in the groups of mice shown in (A). (**C**) Luminescence measurements of Nalm6$^{CD19KO\text{-}CD20}$ cells 14 days after tumor cell engraftment. Data are presented as box plots, with dots indicating measurement from one mouse and lines indicating the median values. In groups without antibody, n = 4 for no T cells, and n = 5 for CAR and CAR$^{FcR}$; in groups with antibody, n = 5 for no T cells and CAR, and n = 7 for CAR$^{FcR}$. P values were calculated by Mann-Whitney test. (**D**) Kaplan-Meier curves show leukemic-free survival; they were analyzed by the log-rank (Mantel-Cox) test. * P < 0.05; **P < 0.01; ***P < 0.001; n.s., not significant.

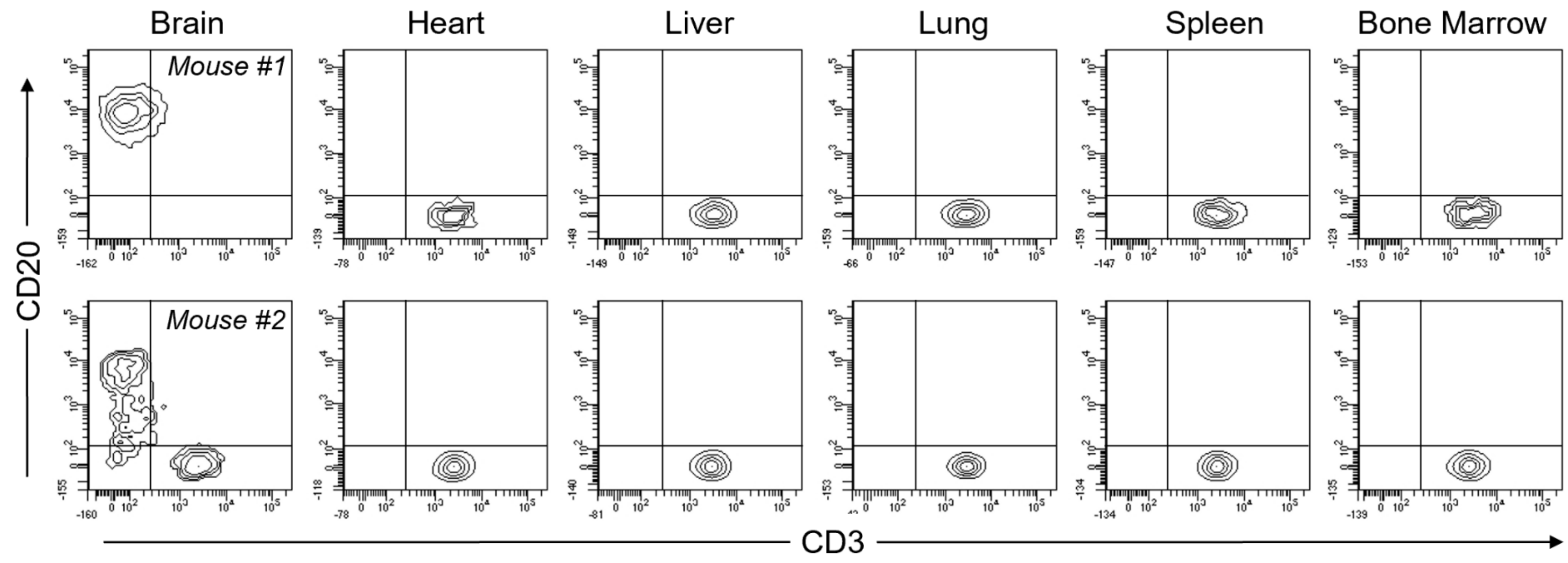


**Figure S6. Flow cytometric analysis of organs collected from mice with recurrent disease after treatment with anti-CD19 CAR$^{FcR}$ and rituximab.** Flow cytometry plots show Nalm6$^{CD19KO\text{-}CD20}$ cells (CD20$^{+}$CD3$^{-}$) and T cells (CD3$^{+}$) in the organs harvested from two mice with relapsed disease after treatment with anti-CD19 CAR$^{FcR}$-T cells and rituximab. Data for one additional mouse is shown in Fig. 4E.

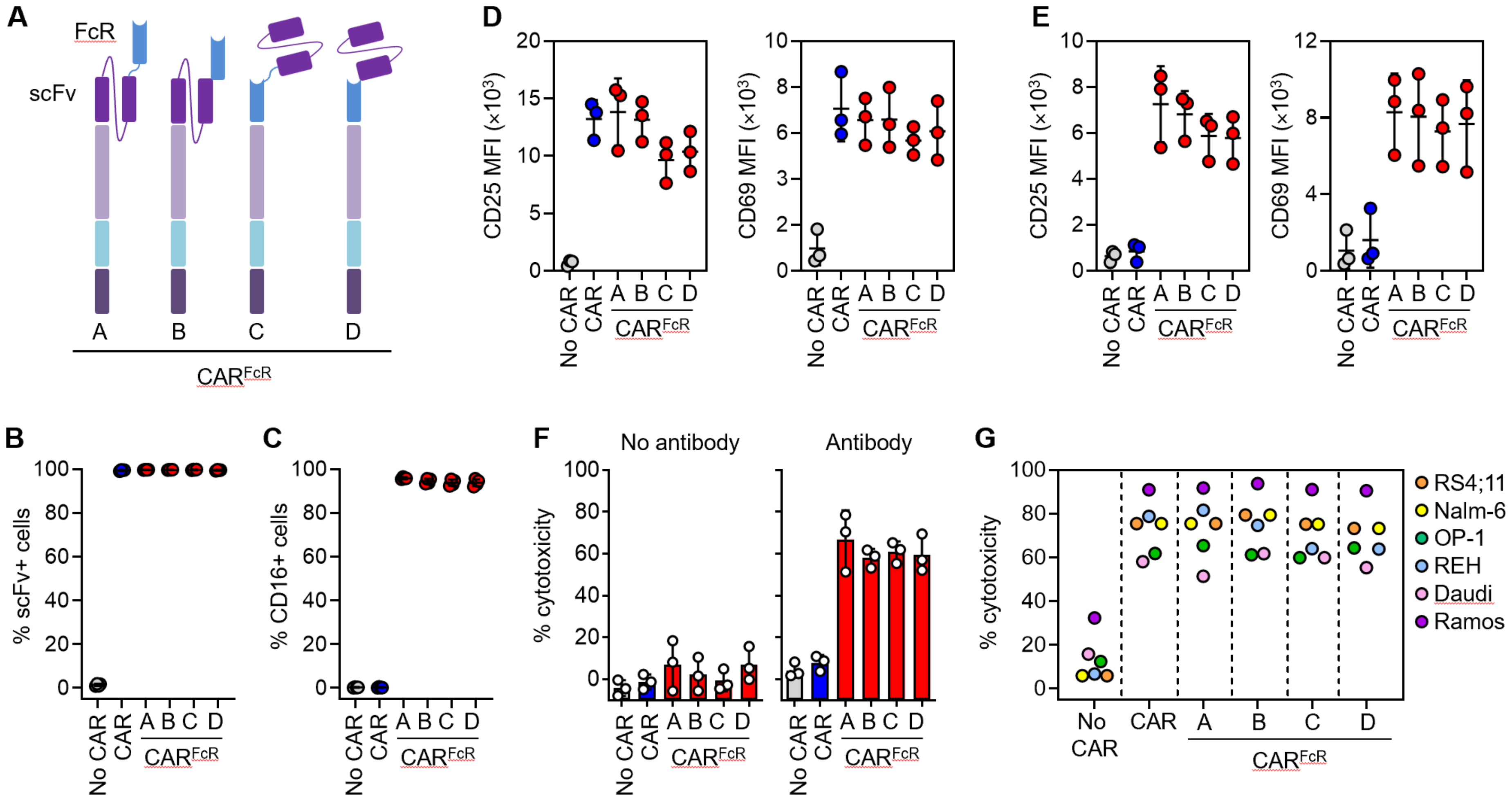


**Figure S7. Expression and function of anti-CD19 CAR$^{FcR}$ variants.** (**A**) Schematic representation of anti-CD19 CAR$^{FcR}$ variants. In CAR$^{FcR}$-A (the standard CAR$^{FcR}$ design), the CD16 FcR with V158 polymorphism is fused to the N-terminus of anti-CD19 scFv via a G4S1 linker, whereas this linker is absent in CAR$^{FcR}$-B. In CAR$^{FcR}$-C and CAR$^{FcR}$-D, the domain orientation is reversed, with a linker present only in CAR$^{FcR}$-C. All constructs have the same components of CAR$^{FcR}$-A shown in Fig. 1A (CD8 hinge and transmembrane domain, and 4-1BB

(CD137) and CD3ζ. (**B**) Percentage of $GFP^+$ cells expressing anti-CD19 scFv. (**C**) Percentage of $GFP^+$ cells expressing CD16. (**D** and **E**) CD25 and CD69 expression in T cells transduced with anti-CD19 $CAR^{FcR}$ variants after 24-hour stimulation with RS4;11 cells (D) or antibody cross-linking (E). (**F**) Cytotoxicity of T cells expressing anti-CD19 $CAR^{FcR}$ variants against $HER2^+$ SKBR3 at a 4:1 E:T ratio with trastuzumab in 4-hour assays. (**G**) Cytotoxicity of T cells expressing anti-CD19 $CAR^{FcR}$ variants against $CD19^+$ ALL and lymphoma cell lines at a 1:1 E:T ratio in 4-hour assays. Each symbol indicate the mean of three biological replicates with each cell line. Data for individual cell lines is shown in Fig. S8. Data for (B) to (F) are mean ± SD; n = 4 biological replicates for (B) and (C); n = 3 biological replicates for (D) to (F).

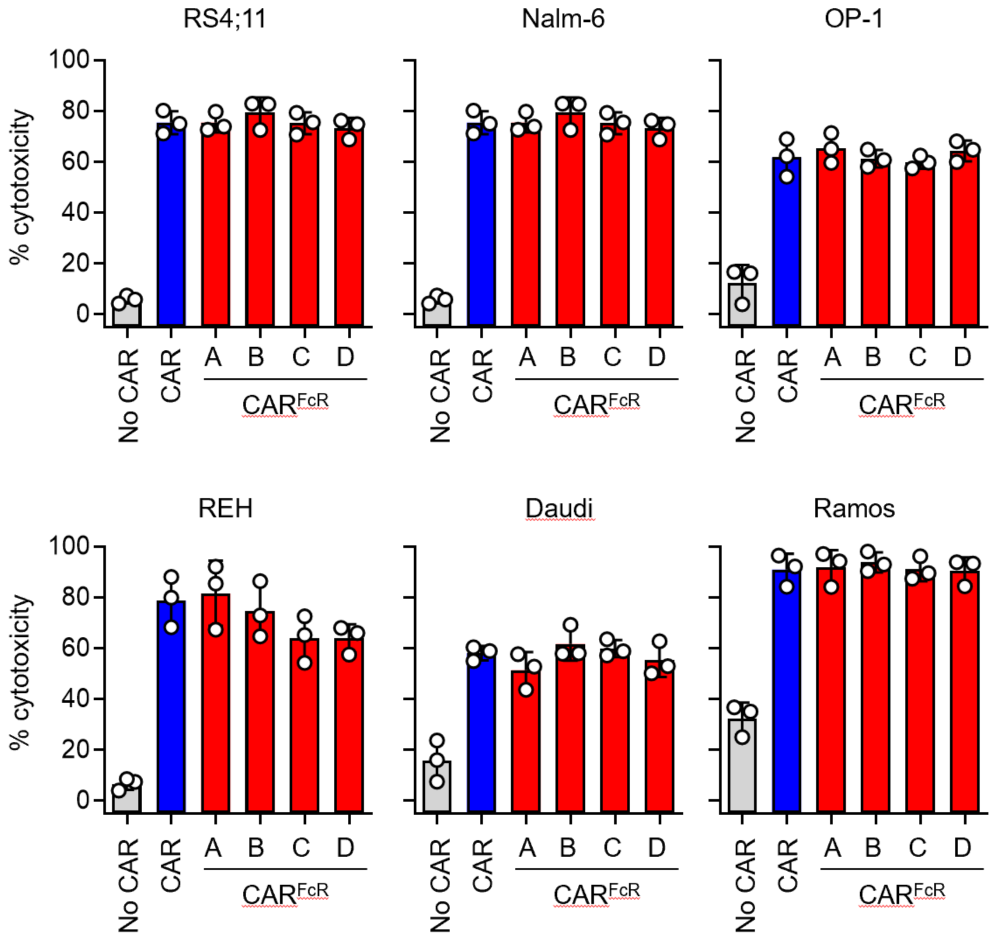


**Figure S8. Cytotoxicity of T cells expressing anti-CD19 CAR$^{FcR}$ variants against CD19$^{+}$ cell lines.** Cytotoxicity of T cells transduced with GFP only (no CAR), anti-CD19 CAR or anti-CD19 CAR$^{FcR}$ variants against CD19$^{+}$ ALL cell lines RS4;11, Nalm-6, OP-1, and REH, and lymphoma cell lines Daudi and Ramos at a 1:1 E:T ratio in 4-hour assays. Data shown are mean ± SD; n = 3 biological replicates.

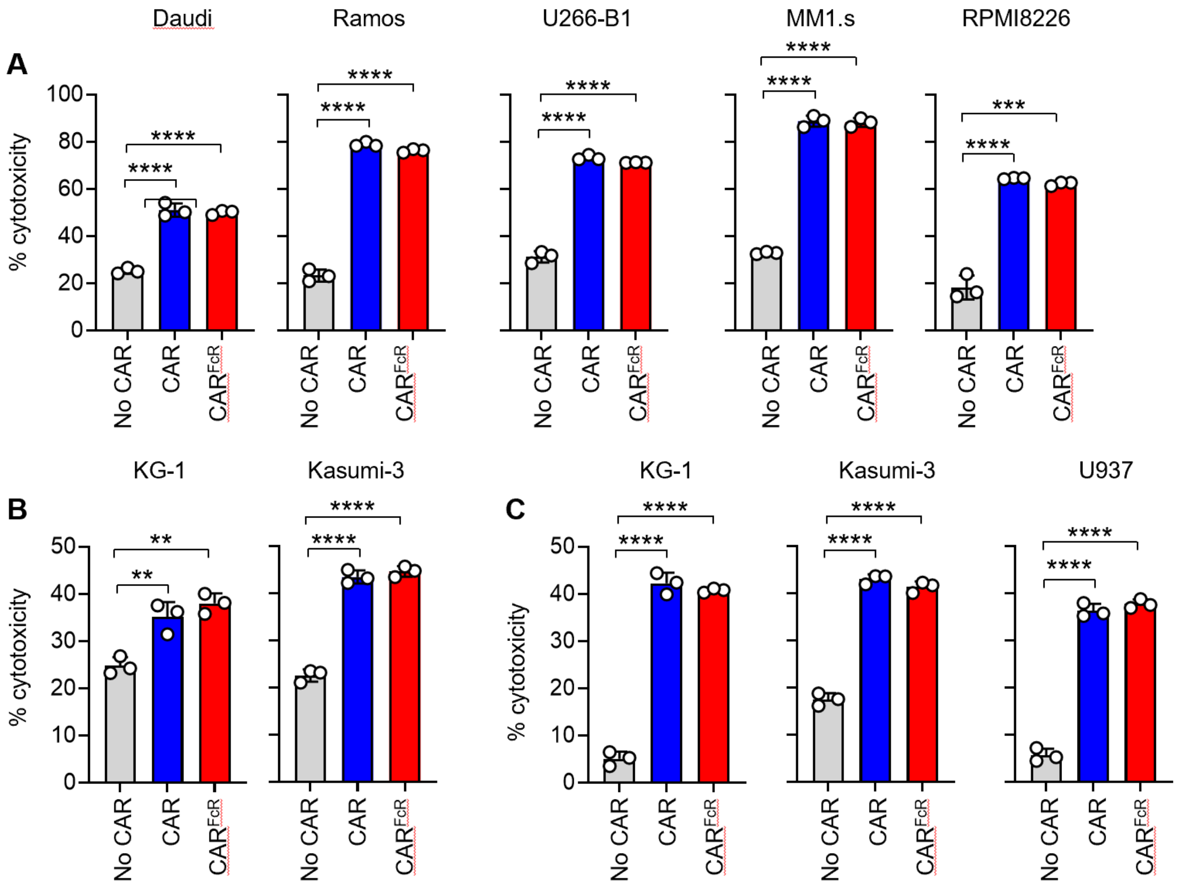


**Figure S9. Cytotoxicity of T cells expressing CAR$^{FcR}$ with anti-BCMA, CD123 or CD33 scFvs.** (**A** to **C**) Cytotoxicity of T cells expressing anti-BCMA (A), anti-CD123 (B), or anti-CD33 (C) CAR or CAR$^{FcR}$ against their respective target cells at a 1:1 E:T ratio in 4-hour assays. Data shown are mean ± SD, with P values determined by unpaired t tests; n = 3 technical replicates. **P < 0.01; ***P < 0.001; ****P < 0.0001.
